# Practical considerations for improved reliability and precision during compound specific analysis of δ^15^N in amino acids using a single combined oxidation-reduction reactor

**DOI:** 10.1101/638098

**Authors:** Philip M. Riekenberg, Marcel van der Meer, Stefan Schouten

## Abstract

**RATIONALE:** There has been increased interest in the analysis for δ^15^N in amino acids to gain simultaneous insight into both trophic relationships and source producers within ecosystems. New developments in gas chromatography combustion isotope ratio mass spectrometry equipment has led to variable outcomes in performance due to limited information about best practices for new systems.

**METHODS:** Precision for δ^15^N in amino acids using the single combined oxidation-reduction reactor is improved across a sequence of analyses if the reactor is oxidized for a substantial period (2 h), immediately followed with a conditioning run of alkanes prior to analysis for N, and the liquid N_2_ CO_2_ trap is left immersed throughout. A five point calibration curve using amino acids with a range of δ^15^N values from −2.4‰ to +61.5‰ was used in combination with a 13 amino acid mixture to correct for offsets during derivatization.

**RESULTS:** Combining the improved setup with normalization techniques using both internal and external standards allows for a reliable throughput of ~25 samples per week. It allowed for a reproducible level of error of <±0.5‰ within standards repeated 10 times across each sequence and a sample error of (±0.18‰), which is lower than analytical error typically associated with δ^15^N-amino acid analysis (±1‰).

**CONCLUSIONS:** A few practical considerations regarding oxidation and conditioning of the combustion reactor allow for increased sequence capacity with the single combined oxidation-reduction reactor. These considerations combined with normalization techniques result in a higher throughput and reduced analytical error during analysis of δ^15^N in amino acids.

## Introduction

There is increasing interest from ecologists in the information about trophic position and source isotope values that are obtained from analysis of δ^15^N in individual amino acids (AA) (Chikaraishi et al. 2007; McMahon and McCarthy 2016; Ohkouchi et al. 2017; Whiteman et al. 2019). Analysis of the differences in δ^15^N between protein-derived amino acids has allowed for the classification of certain amino acids as source and trophic amino acids. Source amino acids are not heavily fractionated during metabolism due to minimal exchange with the metabolic amino acid pool (O’Connell 2017). Due to this, source amino acids reflect the δ^15^N of underlying biogeochemical N sources within the environment that the animal resided as the tissue was formed (Lorrain et al. 2015; Vander Zanden et al. 2013; Vane et al. 2018). Trophic amino acids undergo considerable fractionation as they are metabolized via transamination (Braun et al. 2014; O’Connell 2017) that results in a stepwise enrichment of δ^15^N values that correspond to the trophic level of the animal (Bradley et al. 2015; Chikaraishi et al. 2007; Nielsen et al. 2015). The difference in δ^15^N value between source and trophic amino acids can thus be used to calculate trophic position (TP), a baseline-normalized estimate of the trophic level of that individual within the local food web. It is common to use either the δ^15^N value for glutamic acid (Glu) as a trophic amino acid and phenylalanine (Phe) as a source amino acid (Chikaraishi et al. 2007; Ohkouchi et al. 2017; Vander Zanden et al. 2013) or a combination of several trophic and source AAs. Differences between trophic and source AAs are larger in animals that are secondary and tertiary predators in food webs (Chikaraishi et al. 2007; Nielsen et al. 2015), allowing for quantification of a trophic discrimination factor (TDF) and β, the isotopic differences (‰) between trophic positions within an ecosystem for consumers and primary producers, respectively. Error values associated with TDF and *β* (+1.5‰, (Mcmahon and McCarthy 2016) are larger than the currently accepted measurement error (±1 ‰) for δ^15^N-AA analysis (Yarnes and Herszage 2017).

Analysis of δ^15^N of amino acids via gas chromatography combustion isotope ratio mass spectrometry (GC-C-irMS) requires three steps to isolate and prepare samples for analysis: 1) release of free and bound amino acids from a tissue, 2) isolation of AAs from contaminating matrix materials, and 3) derivatization to make the AAs amenable to interaction with the chromatography column for separation (Ohkouchi et al. 2017). Acid hydrolysis of proteinaceous tissue generally occurs through heating sample material in a strong acid to 100° C for 1-10 h, thereby breaking peptide bonds and releasing the individual AAs from the proteins contained in sample material. Isolation of AAs from interfering sample material is performed if samples contain a complex matrix (e.g. bone, sediment, carbonates) that can interfere with analysis and is commonly done using cation-exchange chromatography (Metges and Petzke 1997; Vane et al. 2018). Use of a cation exchange resin generally results in minimal fractionation of AAs and has sufficient recovery of sample material to allow analysis (Takano et al. 2010). Prior to analysis, derivatization of the AAs is required to reduce polarity and increase volatilization. Derivatization allows for increased chromatographic separation and several derivatization agents can be used e.g. triflouroacetyl-isopropyl ester (McCarthy et al. 1998); pivaloyl-isopropyl ester (Chikaraishi et al. 2007); or methoxycarbonyl AA ester (Walsh et al. 2014). The derivatization used depends on application, and should be explored with consideration towards target amino acids, whether *δ*^13^C or δ^15^N values are desired, the relative stability of derivatized compounds, and introduction of halogenated compounds into the GC-C-irMS (Ohkouchi et al. 2017; Yarnes and Herszage 2017).

Once the clean, derivatized AAs are obtained, they are analyzed via gas chromatography-combustion isotope ratio mass spectrometry (GC-C-irMS) through column separation of the AAs’ followed by combustion and then reduction. A typical combustion/reduction GC interface for the irMS utilizes two reactors, first a Cu/NiO/Pt oxidation reactor heated to 950°C that allows for combustion followed by a Cu reduction reactor (650°C) primarily to reduce NO_x_ products produced during combustion. Common problems encountered during analysis of *δ*^15^N values for AAs via the two separate reactor system include considerable overloading of AA carbon on the GC column as well as the combustion reactor, incomplete combustion, incomplete reduction, leakage (high background N_2_), and issues with standardization to correct for drift occurring across runs. Overloading of AA carbon is expected as there is ~9× more carbon contained in AAs than nitrogen. Therefore, considerable volume of AAs must be loaded onto the GC in order to realize adequate N sample peaks for analysis. Consistent overloading of carbon leads to relatively quick replacement of pre-columns and degeneration/clogging of the combustion reactor and requires more frequent oxidation (and ultimately replacement) of the reactor to maintain combustion capacity. Incomplete reduction results in the appearance of considerable mass 30 (m/z 30, Fig. 1A), derived from the formation of NO_x_, and occurs regularly after long oxidations (60 min) as well as short (e.g. <2 min) routinely used to recover combustion capacity. Leakage within the GC is also a routine problem due to leaks that develop during repeated heating cycles and result in regular downtime. Proper standardization across daily sequences is important to correct for drift in *δ*^15^N values that can occur as the oxidation state of the combustion reactor changes across the day.

**Figure 1:**
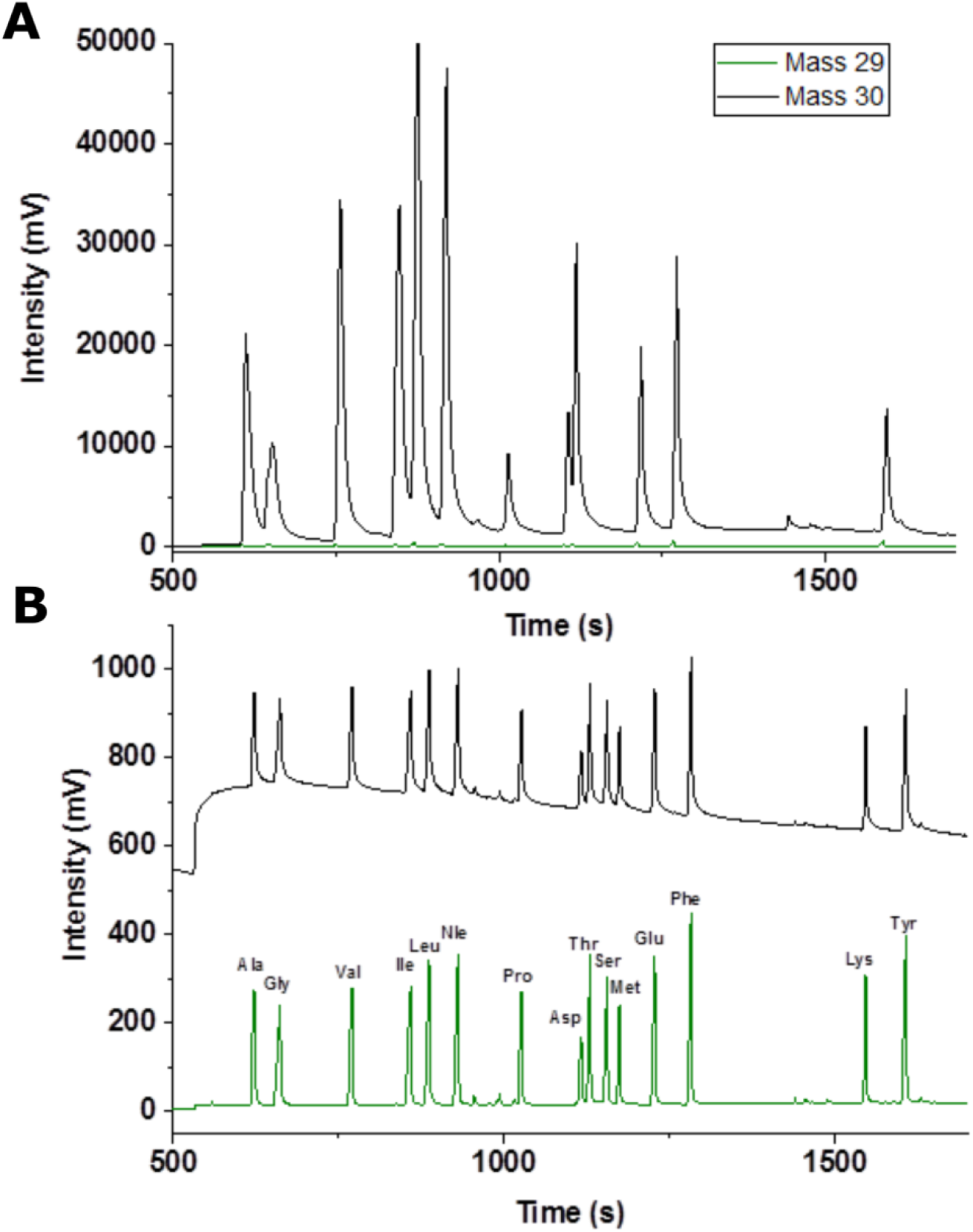
Chromatograms for amino acid standards run immediately after oxidation of the combustion reactor on A) the GC combustion III interface and B) the single combined oxidation-reduction reactor and Isolink II interface. Note the difference in y axis scale between the two plots. For acronyms of amino acids see main text.

The combined effect of these difficulties results in a low sample throughput while ecological studies examining stable isotopes often involve large sample sizes for a number of species within food webs (e.g. Christianen et al., 2017). Recently, a combined oxidation/reduction reactor has become available containing a nickel oxide tube with CuO/NiO/Pt wires (Yarnes and Herszage 2017). Here we discuss the optimization of the combined oxidation/reduction reactor system for compound specific δ^15^N isotopes and incorporating a few practical changes into the daily sequence of analysis. With these changes, we were able to considerably increase sample throughput while achieving increased accuracy and precision during analysis of δ^15^N in individual AAs.

## EXPERIMENTAL

### Reagents and reference materials

Individual amino acids (L-form, >98% purity) were purchased (Sigma Aldrich and Arndt Schimmelmann, Indiana University, respectively) and standard mixtures were created in house. A mix of five amino acids with a known large δ^15^N range (−2.4‰ to 61.5‰; Supplementary table 1) and the internal reference standard L-norleucine (Nle) was used for scale normalization and will be referred to as the ‘scaling AA mix’. This mixture consisted of approximately equimolar quantities of alanine (Ala), glutamic acid (Glu), glycine (Gly), phenylalanine (Phe), Nle, and valine (Val) prepared in 0.1M HCl to a concentration of ~44 mM. A second mixture of 13 amino acids and the internal reference standard (Nle) was used as a reference standard to calculate offsets occurring due to derivatization of AAs and will be referred to as the ‘offset AA mix’. This standard consisted of approximately equimolar quantities of Ala, Aspartic acid (Asp), Glu, Gly, leucine (Leu), isoleucine (Ile), methionine (Met), Phe, threonine (Thr), tyrosine (Tyr), serine (Ser) and Val prepared in a 0.1 M HCl to a concentration of ~8.8 mM. The δ^15^N value for each amino acid used in this mixture was established via direct measurements using an elemental analyzer isotope ratio mass spectrometer (EA-irMS) (Supplementary table 1). An internal reference spike of L-Nle for inclusion into samples after acid hydrolysis was prepared at 44.8 mM in 0.1M HCl. All three solutions were stored in the dark at −20°C.

The solvents and reagents for acid hydrolysis and derivatization included acetyl chloride (>99%, Fluka), bi-distilled water, hydrochloric acid (VWR International), magnesium sulfate (99.5% min, anhydrous; Alfa Aesar), trimethylacetyl chloride (>98%; Alfa Aesar) and HPLC-grade dichloromethane (DCM), ethyl acetate, hexane, isopropanol, and methanol (Promochem; all solvents).

### Sample Materials

Sample materials analyzed included muscle tissue from brown shrimp (*Crangon crangon*), European bass (*Discentrarchus labrax*), mussel (*Mytilus edulis*), European plaice (*Pleuronectes platessa*), and harbour porpoise (*Phocoena phocoena*) collected from the Wadden Sea. Whole body tissue excluding the gut was utilized for ragworm (*Hediste diversicolor*) and plankton (<200 µm mesh) was sampled from the sublittoral zone at Marsdiep, Netherlands (N 53° 0’ 13”, E 4° 46’ 26”) and predominantly consisted of large microalgae. Materials were freeze dried for 48 h and homogenized prior to analysis. These samples reflect a cross section of the Wadden Sea food web with representatives from several trophic positions including primary producer, primary consumer, apex predators and an omnivore (Christianen et al., 2017 and references cited therein).

### Acid Hydrolysis

About 5 mg of freeze dried, ground, and homogenized tissue was placed in a 1 ml reaction vial (Supelco PN: 33293). Vial threads were covered in PTFE tape, 0.5 ml of 6 M HCl was added to the vial, and then vials were capped with a mininert valve cap (15 mm, Supelco PN: 33301). Reaction vials were then placed with valve cap open on a dry heating block set at 100°C with pH indicator paper on top. Vials were closed once the indicator paper showed acidity indicating that oxygen had been purged from the vial. Vials were heated at 100°C overnight (~10-12 h) to allow acid hydrolysis to break down proteins into individual amino acids. After hydrolysis samples were allowed to cool, the acidified water was transferred to a centrifuge filter (GHP Nanosep, 0.45 µm, Pall) and centrifuged (15 s, 10,000 RPM). After filtration, the liquid was transferred to a clean reaction vial, had 0.3 ml of *n*-hexane/DCM (3/2, v/v) added, was capped, vigorously shaken by hand (10 s), and the water and organic solvent were allowed to separate. The top organic solvent layer was pipetted off and discarded and this step was repeated 2× to further clean the extract. 0.2 ml of methanol was added to the remaining hydrolysate (bottom layer).

### Derivatization

Amino acid hydrolysates and amino acid standard mixtures containing water and methanol were dried under N_2_ at 40°C after spiking with 40 uL of NLe (1.8 µmol N). Derivatization followed the procedures according to Svensson et al. (2016) for amino acids with modifications. The amino acids were isopropylated by adding 0.3 ml of a mixture of isopropanol and acetyl chloride (1/4, v/v; prepared by mixing in acetyl chloride dropwise using minimal total volumes of both reagents) and heating at 100°C for 2 h. During initial heating, the valves on the reaction vial caps were not sealed until escaping gasses reacted with pH paper on top of vent to ensure complete flushing of oxygen from the vial. After heating vials were allowed to cool to room temperature. The mixture was then evaporated to dryness under a gentle stream of N_2_ while heated to 40°C. Subsequently 0.25 ml of dichloromethane (DCM) was added and allowed to evaporate to dryness (2×, at 40°C) to remove any remaining reagents. AA *i*-propyl esters were then acylated with a 0.3 ml of a mixture of trimethylacetyl chloride and DCM (1/4,v/v) by heating to 100°C for 1 h to form N-pivaloyl-amino acid-i-propyl esters (NPiP). After heating vials were allowed to cool to room temperature. The reagent was then evaporated to dryness under a gentle stream of N_2_ while heated to 40°C. Subsequently, 0.25 ml of DCM was added and allowed to evaporate to dryness (2×, at 40°C) to remove any remaining reagent. Once dry, 0.2 ml of bi-distilled water and 0.5 ml of *n*-hexane/DCM (3/2, v/v) were added to each reaction vial and shaken for 10 s. The two phases were allowed to separate and the top organic solvent phase was transferred onto a column containing MgSO_4_ (column and MgSO_4_ previously combusted at 450°C) to ensure complete removal of water from the sample. The addition and removal of 0.5 ml of *n*-hexane/DCM to the reaction vial was repeated 2× and then the MgSO_4_ column was rinsed with 0.25 ml of *n*-hexane/DCM to ensure complete removal of the sample from the column. The organic solvent fraction was then allowed to evaporate to dryness under a gentle stream of N_2_ while heated to 40°C. The final evaporation step was closely monitored as the derivatized amino acids are volatile and can be lost at this stage. Ethyl acetate (water removed by addition of 1 g MgSO_4_/100 ml and degassed; by sonicating for 15 min) was then added to the sample and the sample was then stored at −20°C prior to analysis on GC-C-irMS. Prepared NPiP extracts have an expected storage capacity of <12 weeks (Corr et al. 2007) but all samples in this study were run within a week of being derivatized.

### GC-C-irMS analysis

Analyses were performed using a Thermo Trace 1310 gas chromatograph connected to a Delta V Advantage irMS via a GC IsoLink II combustion interface (Thermo Fisher Scientific). The combined oxidation-reduction reactor was a NiO tube containing CuO/NiO/Pt wires within an Alox tube continuously heated to 1000°C during operation. Oxidation capacity of the reactor was insured prior to each sequence by oxidation for 2 h, followed by a backflush period (30 min) to clear any unwanted oxidation products (NO_x_ monitored on m/z 30). The combustion reactor was conditioned after oxidation by a run of alkanes (~equimolar quantities of C_17_, pristane, C_18_, C_20_, C_28_, C_30_, and C_32_; ~25 ng ul^−1^) and oxidation capacity was maintained throughout the sequence by a seed oxidation (12 s) prior to each run during the sequence. Flow rates (GC + backflush (BF), GC, and GC + BF + oxygen) as measured via internal flow meter and free O_2_ (monitored at m/z 32) were closely monitored between sequences as indicators of declining reactor oxidation capacity or potential blockage across the lifetime of the reactor. Using this oxidation program, m/z 30 was held to acceptable levels that did not interfere significantly with δ^15^N across the run (standard deviation <±0.5‰ for 13 AAs across 10 runs of the internal AA standard mix in each sequence) and remained relatively low across sequences (m/z 30 < 700 mV, ratio of m/z 30 / m/z 28 ~5%).

The NPiP amino acid derivatives were injected at ~75°C (PTV with on-column liner for on-column injection; glass, S+H Analytic, Germany) and separated on a DB-5MS column (60 m × 0.32 mm o.d. × 0.5 µm film thickness; Agilent Technologies) at a constant flow rate of 1.6 mL min^−1^ under the temperature program: 70°C for 1 min; ramp to 165°C at 50°C min^−1^; ramp to 185°C at 2°C min^−1^; ramp to 300°C at 10°C min^−1^ and held at 300°C for 7 min. This analysis occurs with a ~1.5 m pre-column (FS deactivated, 0.53 mm) inserted before the column to protect the analytical column. Bad parts (dark spots and burned) of this pre-column can then be routinely cut, mostly from within the injector, or it can be completely replaced. Every third sequence an ~15 cm section of the pre-column is cut off and discarded in order to counter the extensive tailing of peaks that occurs as material builds up in the pre-column within the injector. As the pre-column gets shorter, this approach has diminishing returns and tailing becomes increasingly present, resulting in increasingly noisy standard deviations due to baseline disturbance from peak tails (Ohkouchi et al. 2017). Ultimately, replacement of the entire pre-column serves to resolve this tailing. This strategy allowed for preservation of the gas tight press-fit connecting the pre-column and column for longer between pre-column changes and helped to reduce downtime.

To prevent interference with the measurement of N_2_ in the source, CO_2_ was removed post combustion by a liquid N_2_ trap. CO_2_ was emptied from the trap prior to the 2 h oxidation and the capillary remained outside of the trap until after conditioning with *n*-alkanes was finished. As long as overloading of sample concentration did not occur, there was sufficient capacity within the CO_2_ trap to hold CO_2_ across the entire sequence of 25 runs without emptying. To avoid overloading of the column and CO_2_ trap, all samples were first run on a GC-FID to identify the relative amount of C contained in the samples in order to target sample dilutions prior to analysis on GC-C-irMS.

#### Data Analysis and Normalization

Underivatized individual amino acids were analyzed by EA-irMS for δ^15^N values of the individual amino acids used for normalization during GC-C-irMS analysis (Known δ^15^NAA; Supplementary table 1). Multiple analyses of each amino acid were made on a Flash 2000 elemental analyzer connected to a Delta V Advantage irMS via a Conflo IV interface and were calibrated using the secondary reference materials acetanilide #1 and urea #2 (δ^15^N of 1.18 ± 0.02 and 20.17 ± 0.06, respectively) which were calibrated on IAEA-N-1 and IAEA-N-2 (Schimmelmann et al. 2009). Precision for this analysis is ±0.1‰.

All derivatized samples were analyzed in duplicate by GC-C-irMS. Normalization and scaling for samples and standards followed the calculations presented in Yarnes and Herszage (2017) and were applied to each sequence. This method uses a set of 3 corrections to normalize the data produced through use of 1) an internal reference spike (Nle) included in all samples and standard mixtures and that undergoes the same derivatization, combustion, and reduction process, 2) an amino acid mixture used to calculate an offset for each individual amino acid resulting from derivatization, and 3) a scaling standard mixture using amino acids with a large range of δ^15^N values calibrated against international standards. First, all samples and standard mixtures were evaluated against the internal reference spike instead of laboratory reference gas due to the principal of comparable treatment. Subsequently, a corrected value (δ^15^NAA_Off_) that accounted for derivatization was calculated for all 13AA in the reference mixture using the equation:

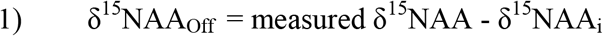

where δ^15^NAA_i_ = measured δ^15^NAA – known δ^15^NAA for each of the amino acids in the 13 AA mixture. δ^15^NAA_Off_ were then normalized to international reference standards by using a linear regression of δ^15^NAA_Off_ values versus the known values of the‘scaling AA mix’ and using the resulting equation to correct the offset adjusted sample values to an internationally calibrated scale across all 5 AAs included in the ‘scaling AA mix’:

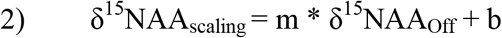

where m is slope and b is the intercept of the regression across all 5 AAs. The materials used for each correction are 1) Nle, 2) ‘offset AA mix, and 3) ‘scaling AA mix’ (δ^15^N ranging from −2.4‰ to +61.5‰; Ala, Glu, Gly, Phe, and Val; Arndt Schimmelmann, Indiana University). δ^15^N values are all presented as ‘per mil’ (‰) relative to atmospheric nitrogen as determined by the International Atomic Energy Agency (IAEA, Vienna, Austria).

### Statistics

Samples and ‘scaling AA mix’ were run in duplicate with a mean ± standard deviation reported for analytical duplicates for δ^15^N_scaling_ values. ‘Offset AA mix’ values are reported as mean ± standard deviation across each sequence (10 replicates). Linear regressions were used to calculate the scaling corrections to internationally calibrated standards (see above). Separation index was calculated as:

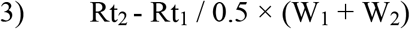

where Rt is retention time and W is width for each of peaks being compared. TP estimates were made using normalized measurements for Glu and Phe within individual samples and calculated as:

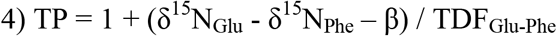

where δ^15^N_Glu_ and δ^15^N_Phe_ are the δ^15^N for Glu and Phe in the sample and TDF and β are set at 7.6‰ and 3.4‰, respectively; (Chikaraishi et al. 2007)

## RESULTS AND DISCUSSION

### Conditioning and monitoring of reactor performance

Our previous method used for measuring δ^15^N of amino acids (e.g. (Svensson et al. 2016) used separate oxidation and reduction reactors in a Thermo Science GC combustion III system. In this method, oxidation by flushing O_2_ was used to improve and maintain combustion capacity only sparingly when peak shape deteriorated (e.g. tailing, reduced peak height). This was due to the considerable increase of m/z 30 intensity (from NOx) and loss of intensity of m/z 28 and 29 (from N_2_) that occurred after this oxidation (Fig. 1A). Recovery of m/z 28 and 29 after oxidation required multiple injections of N-containing compounds (e.g. tri-butyl amine, AA standard mix) to condition both reactors and resulted in considerable loss of analysis time to return to an acceptable measurement baseline. Here, using the combined oxidation and reduction reactor in the Thermo Science IsoLink II system, the time for oxidation and recovery was built into the sequence for each analysis day. In our revised method we included an oxidation period (2 h) prior to sequence start which considerably improved the reliability of the single combined oxidation-reduction reactor during day to day operation. Using the combination of an extended initial oxidation followed with conditioning with a mixture of alkanes and a 12 second seed oxidation before each run, the oxidation capacity of the combustion reactor was considerably improved and behaved more consistently, i.e., prior to this change several reactors clogged close to the weld after minimal use (<2 weeks) when using only seed oxidations (5-10 seconds) between runs to maintain oxidation capacity. The addition of extended oxidation scaled to the N load across the sequence allowed us to analyze 80 samples without problems (in duplicate, 16 sequences with standards presented here) and represents a significant improvement in reliability and laboratory throughput of samples due to practical considerations built into routine analysis procedures.

Besides a 2 h oxidation run, we incorporated into the analytical sequence an ISODAT method that allowed for monitoring of m/z 32 (O_2_) with the backflush turned off between samples and standards. When the oxidation capacity of the reactor was sufficient m/z 32, monitoring O_2_, quickly approached 50 V, but when the oxidation capacity of the reactor was exceeded, m/z 32 would only register 10-30 k mV. This allowed us to monitor exactly within each sequence of runs when/if the oxidation capacity became insufficient and where samples needed to be rerun. Reduction of oxidation capacity routinely resulted in decreased m/z 28 intensities for AAs and increased standard deviation between duplicates within both standards and samples. Furthermore, monitoring carrier gas flow rates prior to each sequence provided additional information about the long term state of the combustion reactor. Declining carrier gas flow rates when the backflush is turned off (GC only) indicated that clogging of the reactor was beginning to occur, although there is some day to day variability in these flow rates (1.56 ± 0.18 ml min^−1^ across the lifetime of the reactor). Directly after analysis, GC flow was >1.4 ml min^−1^ (set at 1.6 ml min^−1^) and m/z 32 is >50000 mV with the backflush turned off. We observed that failure of the reactor was characterized by a gradual increase in σ for δ^15^N in the 13 AA standard mix across a sequence. This gradual failure requires standards to be run throughout the sequence to characterize how the oxidation capacity is maintained towards the end of sequences. Our experience is currently limited to the lifetime of one reactor, and does not represent all possible outcomes for reactor failure (e.g. blockage, leakage).

Another consideration for increased reliability across a daily sequence was to, prior to analysis on GC-C-irMS, analyze samples via GC-FID (DB-5MS, Agilent Technologies, 60 m × 0.32 mm o.d. × 0.5 µm film thickness) in order to ensure that comparable amounts of sample are injected on column for all samples (Ile > 0.3 µg) based on the relative peak areas for sample versus the internal standard spike (Nle, ~0.3 µg, Table 1). This allowed for targeted dilution for both samples and standards prior to on-column injection onto the GC-C-IRMS. Maintenance of comparable analyte loading rates allowed for the CO_2_ trap to remain submerged throughout a sequence after conditioning with alkanes (10 ‘offset AA mix’, 2 ‘scaling AA mix’, and 5 samples in duplicate) eliminating any baseline disturbances resulting from the remnants from emptying the trap between runs. When high concentrations of analyte are loaded on column (e.g. accidental evaporation of solvent in a sample vial) it caused increased CO_2_ background as the CO_2_ trap subsequently became overloaded later in the sequence. Interference caused by overloading can be diagnosed by increasing retention times, considerable drift of the δ^15^N values in standards, and increased error between duplicate analyses.

**Table 1:** Retention times and separation indices for analyzed amino acids for both individual methods as well as a comparison between the two methods. Separation index is calculated for both the amino acids present in each run as well as for the comparable amino acids resulting from each method. Separation index values greater than 1 indicate a difference in peak retention times that is larger than the average width of both peaks. Asp and Thr (1.1) co-eluted at higher standard concentrations, but co-elution was not often observed for peaks with separation index values greater than 1.5.

|  | Svensson et al. 2016 |  |  |  | This study |  |  |  | Comparison |
| --- | --- | --- | --- | --- | --- | --- | --- | --- | --- |
|  | Retention Time (min) | Width | Amplitude 28 | Separation index | Retention Time (min) | Width | Amplitude 28 | Separation index | Separation index |
| **Ala** |  |  |  |  | 10.4 | 12.3 | 655 | 2.7 |  |
| **Gly** | 17.8 | 25.5 | 142 | 24.0 | 11.0 | 11.9 | 293 | 7.1 | 16.9 |
| **Val** |  |  |  |  | 12.9 | 14.4 | 589 | 5.8 |  |
| **Leu** |  |  |  |  | 14.3 | 13.8 | 665 | 1.9 |  |
| **Ile** |  |  |  |  | 14.8 | 16.3 | 513 | 3.0 |  |
| **Nle** | 28.8 | 29.3 | 131 | 50.3 | 15.5 | 13.4 | 749 | 7.0 | 20.2 |
| **Pro** |  |  |  |  | 17.1 | 16.1 | 550 | 7.7 |  |
| **Asp** |  |  |  |  | 18.6 | 10.9 | 488 | 1.1 |  |
| **Thr** |  |  |  |  | 18.8 | 11.3 | 427 | 2.0 |  |
| **Ser** |  |  |  |  | 19.2 | 15.5 | 676 | 1.5 |  |
| **Met** |  |  |  |  | 19.6 | 14.4 | 864 | 4.3 |  |
| **Glu** | 49.5 | 20.1 | 96 | 5.6 | 20.5 | 11.7 | 422 | 3.9 | 3.9 |
| **Phe** | 51.4 | 21.5 | 249 | 26.0 | 21.4 | 13.4 | 536 | 20.8 | 21.2 |
| **Lys** |  |  |  |  | 25.8 | 12.5 | 614 | 4.5 |  |
| **Tyr** | 59.6 | 16.3 | 322 |  | 26.8 | 16.1 | 755 |  |  |

### Normalization

Rather than using solely a single AA mixture to account for both changes in δ^15^N due to derivatization of AAs and long-term stability (Svensson et al. 2016), we followed the approach of Yarnes and Herszage (2017) which uses 3 standards to normalize AAs. These standards are: 1) a spiked AA reference standard (Nle) included in every standard and sample, 2) an AA mixture to account for δ^15^N changes during derivatization ‘offset AA mix’(Fig. 1B), and 3) an AA mixture to scale to calibrated international standards ‘scaling AA mix’ (Table 1). Precision was within ±0.5‰ for both standard mixes run during the analysis period (5 weeks; 16 sequences; ‘offset AA mix’: 160 injections, average σ ± 0.22‰, range ± 0.18-0.25, min-max: Met-Ala; ‘scaling AA mix’: 32 injections, σ ±0.19, range ± 0.1-0.33, min-max: Val-Gly; Fig. 1B & Table 1). This represents an improvement over both precision of our previously utilized standard mix (0.7-1.2‰; (Svensson et al. 2016) as well as the commonly reported precision of ±1‰ for analysis of ^15^N in AAs (Walsh et al. 2014; Yarnes and Herszage 2017). Investigating the effects of each correction individually indicated that the mean difference between values of amino acids measured by EA-irMS and their derivatives prepared via the NPiP pathway (δ^15^NAA_i_) was larger when corrected solely for the spiked internal reference material than for standards that were additionally scale-normalized (1.41±0.18‰ vs −0.88±0.22‰, respectively; Fig. 2). Measured differences between AAs measured via EA and their derivatives measured via GC-C-irMS were larger for some amino acids (derivatization offset AA mix: Gly, Thr, and Ser ~2‰, Fig. 2; Scaling AA mix: Ala ~2.3‰, Fig. 2). Samples that were corrected using this method for normalization had an average precision of 0.18‰ for duplicate measurements (0.01-0.49‰ min-max, 12AAs, 7 samples; Table 2) and have been normalized to internationally calibrated reference materials and are comparable to measurements in other studies that are similarly calibrated.

**Table 2:** δ^15^N values for NPiP derivatives from acid hydrolysis of tissue of different species from the Wadden Sea after normalization using both internal and scaling AA mix. Plankton was material from a bulk water sample retained using a 200 µm mesh size and predominantly consisted of large microalgae. N.D. indicates not determined as the peak heights were below 100 mV and therefore were not suitable for integration.

|  | Brown Shrimp | European Bass | Mussel | European Plaice | Ragworm | Harbour Porpoise | Phytoplankton |
| --- | --- | --- | --- | --- | --- | --- | --- |
|  | *Crangon crangon* | *Dicentrarchus labrax* | *Mytilus edulis* | *Pleuronectes platessa* | *Hediste diversicolor* | *Phocoena phocoena* |  |
| **Ala** | 27.31(0.28) | 30.24(0.23) | 19.94(0.04) | 24.62(0.28) | 16.40(0.26) | 22.83(0.24) | 12.98(0.02) |
| **Asp** | 22.7(0.20) | 26.65(0.26) | 16.19(0.05) | 23.00(0.16) | 15.58(0.01) | 23.36(0.15) | 14.12(0.41) |
| **Glu** | 27.65(0.33) | 30.16(0.03) | 18.26(0.20) | 25.52(0.23) | 18.56(0.10) | 26.00(0.13) | 15.79(0.29) |
| **Gly** | 10.25(0.30) | 9.23(0.39) | 8.19(0.11) | 9.85(0.29) | 8.95(0.10) | 20.46(0.03) | 10.98(0.19) |
| **Ile** | 18.8(0.30) | 26.59(0.11) | 18.00(0.29) | 22.56(0.10) | 15.52(0.08) | 20.85(0.29) | 10.42(0.06) |
| **Leu** | 21.29(0.13) | 29.89(0.25) | 16.73(0.13) | 24.37(0.07) | 15.64(0.13) | 24.96(0.22) | 11.54(0.33) |
| **Met** | 12.47(0.09) | 14.12(0.23) | 7.60(0.43) | 12.48(0.11) | 8.40(0.07) | N.D. | N.D. |
| **Phe** | 10.35(0.01) | 10.06(0.36) | 7.75(0.09) | 10.27(0.35) | 6.80(0.21) | 12.85(0.22) | 10.31(0.02) |
| **Ser** | 9.87(0.15) | 12.49(0.28) | 7.34(0.09) | 10.80(0.06) | 7.23(0.08) | 20.06(0.49) | 5.73(0.06) |
| **Thr** | 5.96(0.10) | (-1.4)(0.42) | 4.00(0.10) | 8.15(0.33) | (-1.96)(0.07) | (-16.79)(0.03) | 6.97(0.15) |
| **Tyr** | 12.76(0.27) | 14.52(0.02) | 9.14(0.07) | 12.27(0.23) | 8.46(0.046) | N.D. | 8.87(0.11) |
| **Val** | 22.89(0.27) | 29.71(0.32) | 18.94(0.07) | 24.74(0.20) | 17.05(0.13) | 26.45(0.42) | 15.18(0.19) |
| **Glu-Phe** | 17.29(0.34) | 20.10(0.33) | 10.52(0.29) | 15.25(0.12) | 11.76(0.10) | 13.15(0.35) | 5.48(0.28) |
| **TP** | 2.8 | 3.2 | 1.9 | 2.6 | 2.1 | 2.3 | 1.3 |

**Figure 2:**
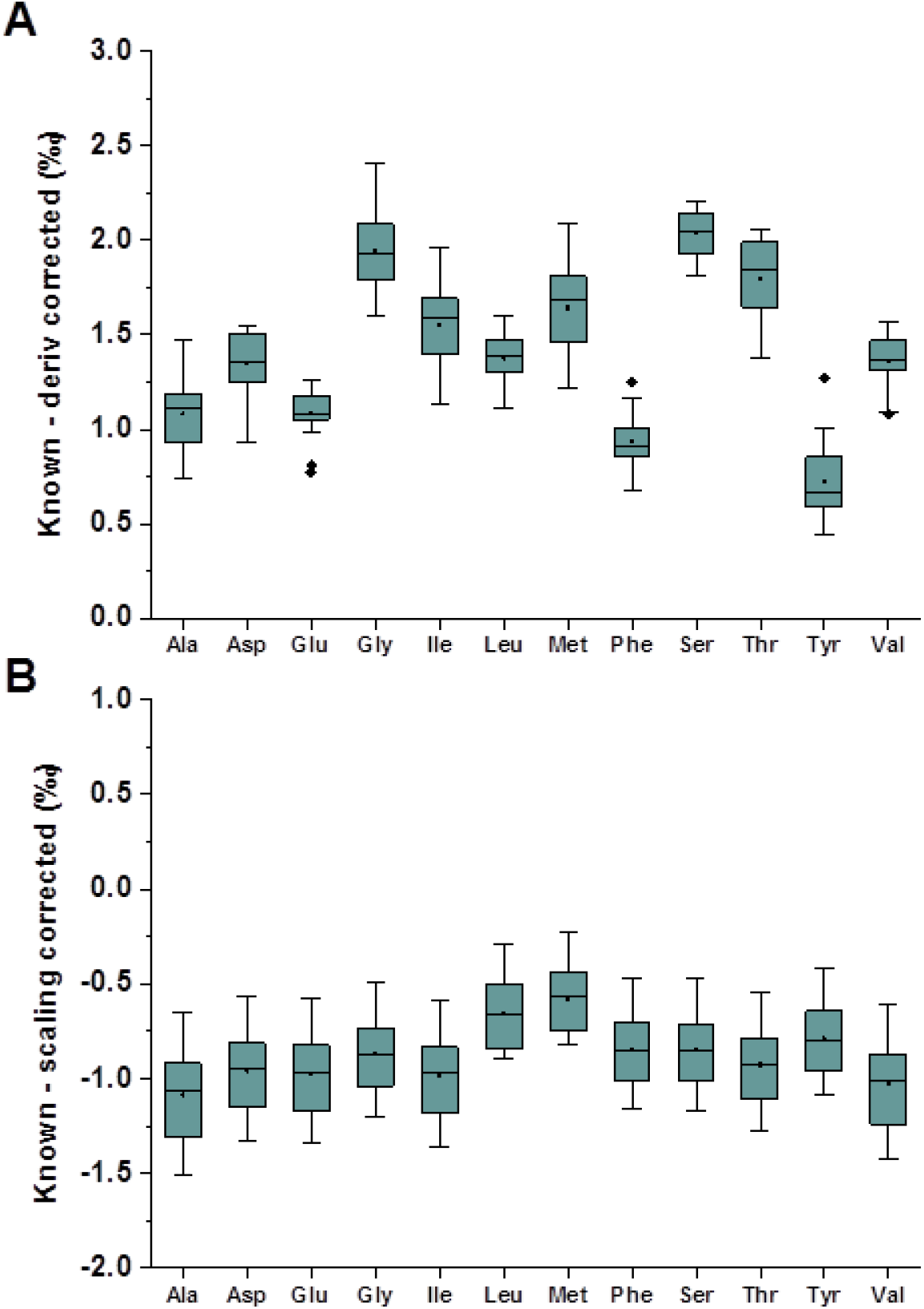
Box plots of measured isotopic offsets for a 12 amino acid standard mixture across 16 daily runs between A) δ^15^N values measured by EA-irMS for free amino acids and by GC-C-irMS corrected for derivatization and B) δ^15^NAA_i_, offsets between known δ^15^NAA values from EA-irMS analysis and δ^15^NAA values measured by GC-C-irMS corrected for derivatization and scaled to internationally calibrated compound specific standards. Within the boxplots, black dots are the mean, lines represent the median, boxes represent the upper and lower quartiles, and whiskers represent the 1.5 quartile ranges. Any black diamonds outside of the whiskers are outliers.

**Figure 3:**
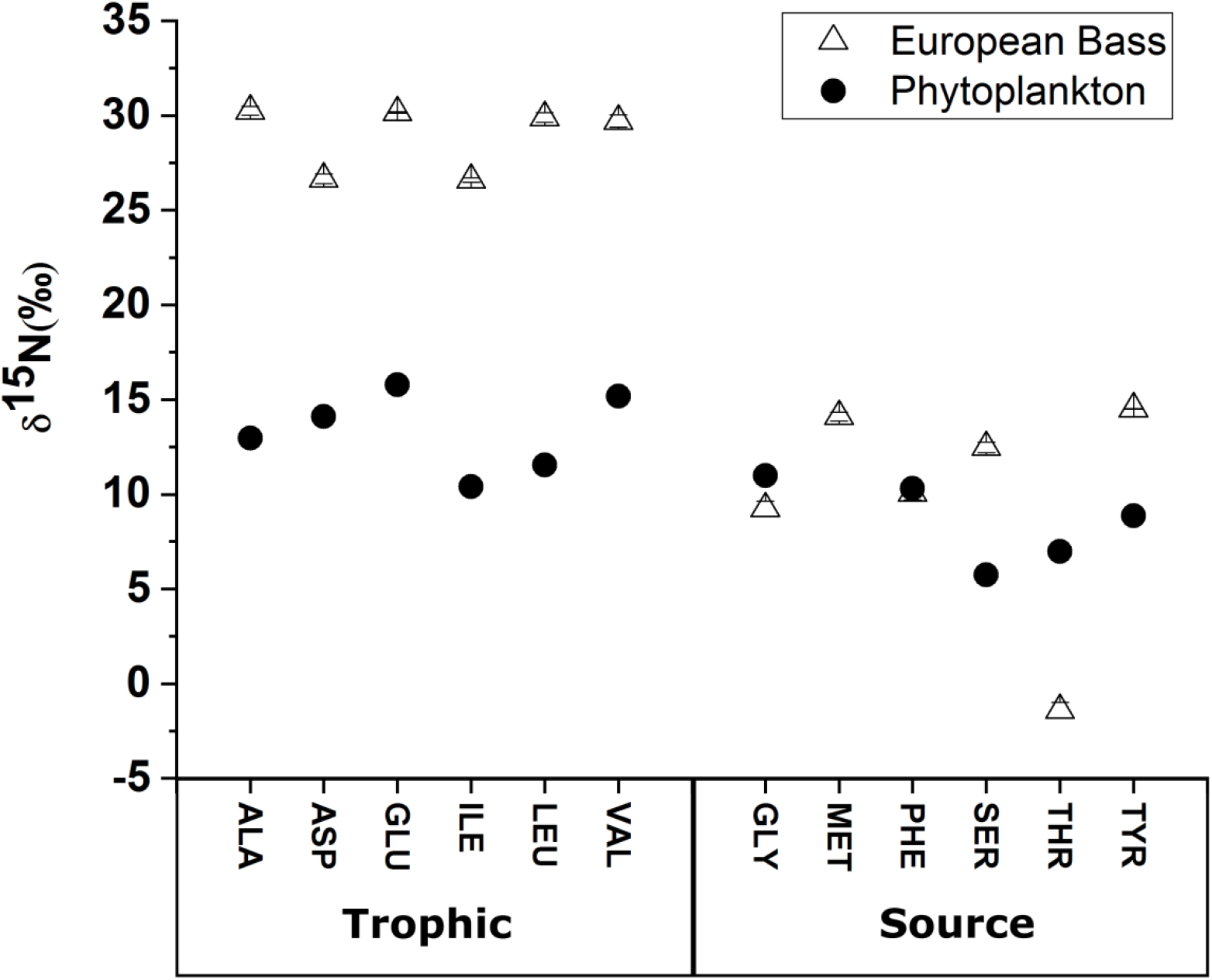
δ^15^N values for 12 amino acids in a fish (European Bass) and phytoplankton (collected at Marsdiep, Netherlands) assigned to “Trophic” and “Source” groups. (Mean ± SD for analytical duplicates, some error bars are too small to be seen)

It should be noted that lysine and proline showed an offset between AAs measured via EA and their derivatives measured via GC-C-irMS that were less consistent than the other 12 reported AAs, and resulted in less consistent determination for δ^15^NAA_i_. The precision for these two AAs fell within the larger ±1 ‰ range during the analysis (Lys 0.78‰, Pro 0.99‰, ± SD; Supplementary figure 1). Decreased precision for these standards may be due to incomplete derivatization during standard preparation or instability in the derivatized product during storage at −20°C. The shifting δ^15^N values for these two amino acids were only discovered through tracking of δ^15^NAA_i_ measured for all AAs across the 16 sequences. This example demonstrates the valuable insight into the long term stability of precision and accuracy during analysis that is gained by tracking the derivatization offset values across sequences. Widespread changes in derivatization offset values across all AAs can indicate that oxidation performance is changing in the reactor and/or deteriorating and that maintenance may be needed.

#### Trophic position

In addition to the “canonical” source and trophic AAs (Phe, Glu), the improved method has also allowed δ^15^N measurements for other source (Gly, Met, Ser, Thr, Tyr) and trophic (Ala, Asp, Ile, Leu, Val) AAs with improved precision (±0.5‰, 1σ; Fig. 1B, Table 1 & 2). Improved precision for non-canonical AAs will allow for further reduction of the variability associated with TDF for both consumers and β as multiple amino acids are increasingly utilized to better constrain variability amongst AA types (Bradley et al. 2015) or alternative trophic AAs (e.g. Pro) are utilized for TP measurements (Vokhshoori et al. 2019). Improvement for estimates of TDF and *β* below the current precision of ±1.5‰ will better reflect metabolic variability occurring within species as measurement error is reduced (Mcmahon et al. 2015; Yarnes and Herszage 2017). Additionally, improved machine reliability and reduced downtime will allow for increased replication for species and better efficiency during method development for difficult materials (e.g. microphytobenthos, detrital and sediment trap organic matter).

To demonstrate the improved procedure we analyzed AAs for 6 species (*Crangon crangon*, brown shrimp; *Dicentrarchus labrax*, European bass; *Hediste diversicolor*, Ragworm; *Mytilus edulis*, blue mussel; *Phocoena phocoena*, harbor porpoise; and *Pleuronectes platessa*, European plaice) and phytoplankton sampled from the Wadden Sea. These samples represent an expected range of animals from the Wadden Sea food web that spans from primary producers to tertiary consumers (Table 2). In addition, we also compared a subset of *M. edulis* samples that were analyzed using both our improved method as well as the previously used (cf. Svensson et al., 2016; n=4; Fig. 5). Both the highest and lowest TPs in this study (*D. labrax* and phytoplankton, respectively; Fig. 3) largely fit expected trends, with elevated δ^15^N values found for the “trophic” AAs in the tertiary consumer when compared to the primary producer. δ^15^N values for the source AAs were generally lower, reflecting the smaller fractionation associated with their processing during trophic transfer, with the notable exception of Thr, which was considerably depleted for the European bass (−1.4‰; Table 2). This negative correlation with elevated trophic position has been previously observed (Bradley et al. 2015; Hare et al. 1991; Mcmahon et al. 2015) and may indicate a strong reverse fractionation associated with trophic transfer of N or be an effect of diet quality or metabolic rate (McMahon and McCarthy 2016). The δ^15^N value for Gly is relatively close to that of Phe (9.2‰ and 10.1‰ respectively) and may reflect a minimal contribution of microbially-degraded material to the diet of European bass (Calleja et al. 2013). This analytical method reliably produces δ^15^N values for non-canonical AAs that should allow for further insight to the metabolic processes occurring during trophic transfer of AA’s between species.

The TP calculated here using Glu and Phe (equation 4) ranged from 1.3 for phytoplankton to 3.2 for *D. labrax* which correlate well with values expected for primary producers to tertiary consumers, respectively (Table 2 & Fig. 4). European plaice, *P. platessa* had a TP of 2.6 likely reflecting a mixture of both predation and detritivory that reflects a bottom feeding lifestyle. The TP for the harbour porpoise *P. phocena* (2.3) was considerably lower than would be expected for a tertiary consumer, but lower TPs have been observed previously for marine mammals (Matthews and Ferguson 2013; Yarnes and Herszage 2017). TP for both *H. diversicolor* and *M. edulis* were similar (2.1 and 2.3 respectively) indicating that both of these species are primary consumers. *C. crangon* had a TP of 2.8, which indicates that it is a tertiary consumer and likely reflects occasional supplemental predation in addition to detritus feeding (Van Der Veer and Bergman 1987).

**Figure 4:**
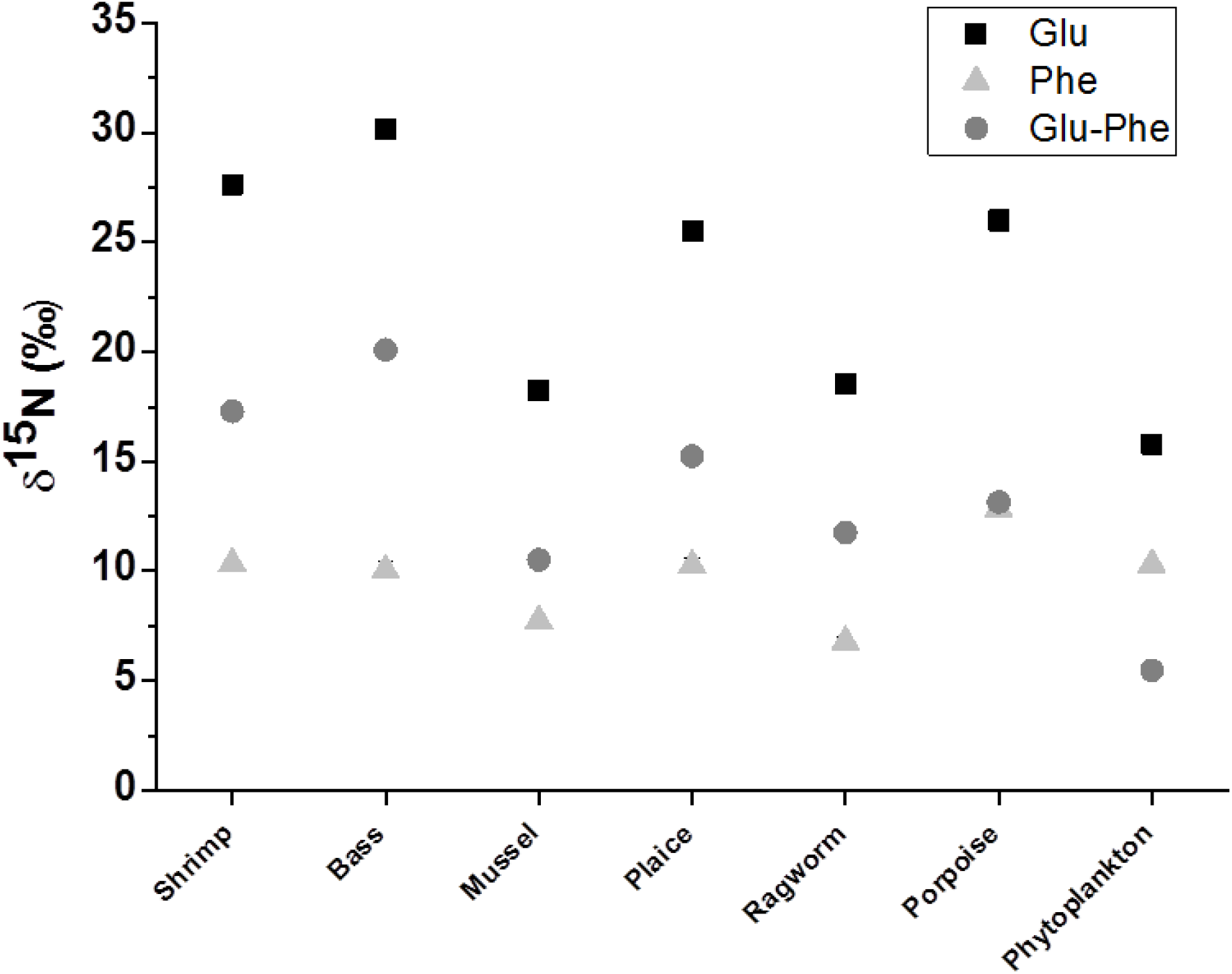
δ^15^N values glutamic acid and phenylalanine for 7 samples as well as the difference that is commonly used in the calculation of trophic position. (Mean ± 0.5SD of analytical duplicates, error bars are too small to be seen)

Comparison between the method presented here and the previous method (Svensson et al. 2016) showed that standard deviations for Glu and Phe, and the resulting TPs were much lower (Glu: 0.7‰ vs 3.3‰, Phe: 0.7‰ vs 4.6‰, Glu: 0.7‰ vs 3.3‰, and TP: 0.1 vs 0.4, respectively; Fig. 5). This decrease was primarily observed due to more consistent standardization techniques that better account for any drift in AA δ^15^N values that may occur across the daily sequences. However, there was considerably less drift and increased stability in δ^15^N values produced using the combined oxidation and reduction reactor and the data were produced at a considerably quicker rate due to increased reliability of the system. Additionally, the δ^15^N values produced have been standardized to international reference standards and will be comparable to similarly standardized measurements. This improvement in analytical precision and more widespread utilization of normalization to international standards will allow for more accurate and precise estimates of TP enabling better comparisons between TPs of animals across ecosystems.

**Figure 5:**
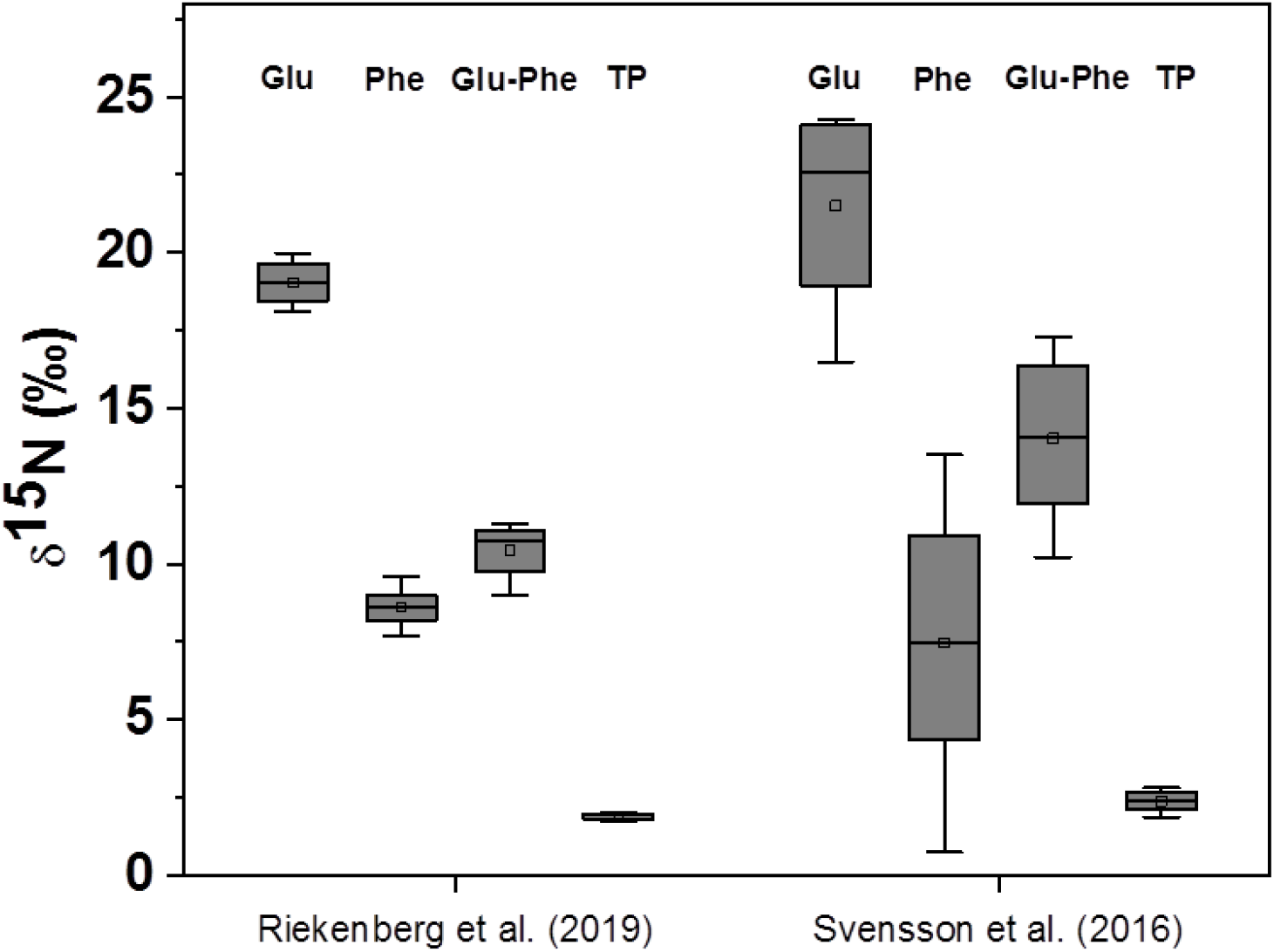
δ^15^N values for glutamic acid (Glu), phenylalanine (Phe), Glu-Phe, and trophic position (TP) for both the method presented here and the previously used method (Svensson et al. 2016). Within the boxplot, open squares represent the mean, lines represent the median, boxes represent the upper and lower quartiles, and whiskers represent the 1.5 quartile ranges. Any black squares outside of the whiskers are outliers.

#### Conclusion

Extended daily oxidation, conditioning with alkanes, use of seed oxidations between runs, and targeted analyte loading rates allowed for increased reliability and precision across the lifetime of a single combined oxidation-reduction reactor used for analysis of δ^15^N in amino acids via GC-C-IRMS. These practical changes were easily built into the daily sequence for amino acid analysis. When these methods are coupled with the use of both internal and scale normalization, the precision realized during ^15^N-AA analysis was <± 0.5‰. CSIA users can utilize these steps to further improve the reliability and stability when using a single combined oxidation-reduction reactor and the IsoLink II.

## Acknowledgements

We thank students from the 2018 NIOZ Marine Master course for providing samples for analysis. We thank Dieter Juchelka and Maria de Castro (Thermo Fisher Scientific, Bremen, Germany) for technical assistance following considerable downtime. We also thank Diane O’Brien (University of Alaska) and Christopher Yarnes (UC Davis) for encouraging output of a relatively user specific method paper during discussion at ISOECOL 2018. Additionally, this work benefited from participation in the ISOECOL 2018 workshop: Approaches to improve the reproducibility of amino acid stable isotope analyses.

---

**Supplementary table 1:** δ^15^N values (‰) for reference standards used for offsets that occurred during derivatization and scale normalization.

|  | 13 AA<br>Standard<br>Mix | 6 AA<br>Scaling<br>Standard<br>Mix |
| --- | --- | --- |
| Ala | -7.4 | 43.25 |
| Asp | -2.4 |  |
| Glu | -2.9 | -2.38 |
| Gly | 1.1 | 20.7 |
| Ile | -3.3 |  |
| Leu | 9.7 |  |
| Met | -2.1 |  |
| Nle | 12.6 | 12.6 |
| Phe | 2.2 | 1.7 |
| Ser | 1.9 |  |
| Thr | -1.1 |  |
| Tyr | 4.3 |  |
| Val | -5.1 | 61.53 |

**Supplementary figure 1:**
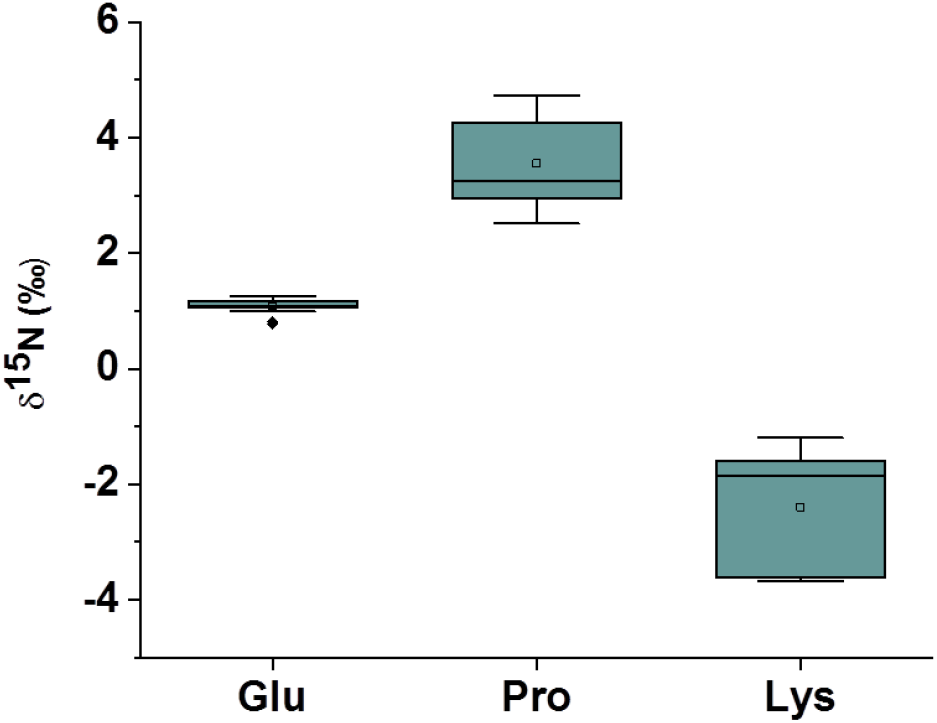
δ^15^N values for lysine and proline compared with glutamic acid across the analysis. Note the larger range for lysine and proline versus glutamic acid (0.78‰, 0.99‰, and 0.13‰, respectively; all ± SD)

## LITERATURE CITED

Bradley, C. J. and others 2015. Trophic position estimates of marine teleosts using amino acid compound specific isotopic analysis. Limnology and Oceanography: Methods 13: 476–493.

Braun, A., A. Vikari, W. Windisch, and K. Auerswald. 2014. Transamination governs nitrogen isotope heterogeneity of amino acids in rats. J Ag Food Chem 62: 8008–8013.

Calleja, M. L., F. Batista, M. Peacock, R. Kudela, and M. D. McCarthy. 2013. Changes in compound specific δ^15^N amino acid signatures and d/l ratios in marine dissolved organic matter induced by heterotrophic bacterial reworking. Mar Chem 149: 32–44.

Chikaraishi, Y., Y. Kashiyama, N. O. Ogawa, H. Kitazato, and N. Ohkouchi. 2007. Metabolic control of nitrogen isotope composition of amino acids in macroalgae and gastropods: Implications for aquatic food web studies. Mar Ecol Prog Ser 342: 85–90.

Christianen, M. J., Middelburg, J. J., Holthuijsen, S. J., Jouta, J., Compton, T. J., Heide, T., Piersma, T., Sinninghe Damsté, J. S., Veer, H. W., Schouten, S. and Olff, H. 2017. Benthic primary producers are key to sustain the Wadden Sea food web: Stable carbon isotope analysis at landscape scale. Ecology 98: 1498–1512.

Corr, L. T., R. Berstan, and R. P. Evershed. 2007. Optimisation of derivatisation procedures for the determination of δ^13^C values of amino acids by gas chromatography/combustion/isotope ratio mass spectrometry. Rapid Communications in Mass Spectrometry 21: 3759–3771.

Hare, P. E., M. L. Fogel, T. W. Stafford Jr, A. D. Mitchell, and T. C. Hoering. 1991. The isotopic composition of carbon and nitrogen in individual amino acids isolated from modern and fossil proteins. J Arch Sci 18: 277–292.

Lorrain, A. and others 2015. Nitrogen isotopic baselines and implications for estimating foraging habitat and trophic position of yellowfin tuna in the Indian and Pacific Oceans. Deep Sea Res II 113: 188–198.

Matthews, C. J. D., and S. H. Ferguson. 2013. Spatial segregation and similar trophic-level diet among eastern Canadian Arctic/north-west Atlantic killer whales inferred from bulk and compound specific isotopic analysis. Journal of the Marine Biological Association of the United Kingdom 94: 1343–1355.

McCarthy, M. D., J. I. Hedges, and R. Bennar. 1998. Major bacterial contribution to marine dissolved organic nitrogen. Science 281: 231–234.

McMahon, K. W., and M. D. McCarthy. 2016. Embracing variability in amino acid δ^15^N fractionation: mechanisms, implications, and applications for trophic ecology. Ecosphere 7: e01511-n/a.

McMahon, K. W., S. R. Thorrold, T. S. Elsdon, and M. D. McCarthy. 2015. Trophic discrimination of nitrogen stable isotopes in amino acids varies with diet quality in a marine fish. Limnology and Oceanography 60: 1076–1087.

Metges, C. C., and K. J. Petzke. 1997. Measurement of15N/14N Isotopic Composition in Individual Plasma Free Amino Acids of Human Adults at Natural Abundance by Gas Chromatography– Combustion Isotope Ratio Mass Spectrometry. Analytical Biochemistry 247: 158–164.

Nielsen, J. M., B. N. Popp, and M. Winder. 2015. Meta-analysis of amino acid stable nitrogen isotope ratios for estimating trophic position in marine organisms. Oecologia 178: 631–642.

O’Connell, T. C. 2017. ‘Trophic’ and ‘source’ amino acids in trophic estimation: a likely metabolic explanation. Oecologia 184: 317–326.

Ohkouchi, N. and others 2017. Advances in the application of amino acid nitrogen isotopic analysis in ecological and biogeochemical studies. Organic Geochemistry 113: 150–174.

Schimmelmann, A., A. Albertino, P. E. Sauer, H. Qi, R. Molinie, and F. Mesnard. 2009. Nicotine, acetanilide and urea multi-level 2H-, 13C- and 15N-abundance reference materials for continuous-flow isotope ratio mass spectrometry. Rap Comm Mass Spectrom 23: 3513–3521.

Svensson, E., S. Schouten, E. C. Hopmans, J. J. Middelburg, and J. S. Sinninghe Damsté. 2016. Factors Controlling the Stable Nitrogen Isotopic Composition (δ^15^N) of Lipids in Marine Animals. PLOS ONE 11: e0146321.

Takano, Y., Kashiyama, Y., Ogawa, N. O., Chikaraishi, Y. and Ohkouchi, N. 2010. Isolation and desalting with cation-exchange chromatography for compound-specific nitrogen isotope analysis of amino acids: application to biogeochemical samples. Rapid Commun Mass Spectrom 24: 2317–2323.

Van Der Veer, H. W., and M. J. Bergman. 1987. Predation by crustaceans on a newly settled 0-group plaice Pleuronectes platessa population in the western Wadden Sea. Mar Ecol Prog Ser 35: 203–215.

Vander Zanden, H. B. and others 2013. Trophic ecology of a green turtle breeding population. Mar Ecol Prog Ser 476: 237–249.

Vane, K., N. J. Wallsgrove, W. Ekau, and B. N. Popp. 2018. Reconstructing lifetime nitrogen baselines and trophic position of Cynoscion acoupa from δ^15^N values of amino acids in otoliths. Mar Ecol Prog Ser 597: 1–11.

Vokhshoori, N. L. and others 2019. Broader foraging range of ancient short-tailed albatross populations into California coastal waters based on bulk tissue and amino acid isotope analysis. Mar Ecol Prog Ser 610: 1–13.

Walsh, R. G., S. He, and C. T. Yarnes. 2014. Compound-specific δ13C and δ15N analysis of amino acids: A rapid, chloroformate-based method for ecological studies. Rap Comm Mass Spectrom 28: 96–108.

Whiteman, J. P., E. A. Elliott Smith, A. C. Besser, and S. D. Newsome. 2019. A Guide to Using Compound-Specific Stable Isotope Analysis to Study the Fates of Molecules in Organisms and Ecosystems. Diversity 11: 8.

Yarnes, C. T., and J. Herszage. 2017. The relative influence of derivatization and normalization procedures on the compound-specific stable isotope analysis of nitrogen in amino acids. Rap Comm Mass Spectrom 31: 693–704.

